# Positive Reinforcement Training Insight from Working Elephants in India

**DOI:** 10.1101/2025.11.15.688661

**Authors:** Kharkar Rutwik, Velho Nandini, Rathod Shradha, Boro Ashok, Pratt Chrissy, Rathod Pooja, Rambia Kime

## Abstract

Working elephants in India are predominantly trained using dominance-based and punitive methods. Research over the past few decades has shown that these methods, when contrasted with positive reinforcement training techniques, can be detrimental to animals’ mental and physical well being. Positive reinforcement based methods have also been shown to be effective in obtaining the voluntary participation of many species of animals in husbandry and routine medical procedures. Our study aims to characterize the stress responses of a group of captive working elephants longitudinally while desensitizing certain procedures and training assistive behaviours. We documented the duration and frequency of stress signals presented by elephants during regular training sessions over a four month period. We found that as sessions progressed, elephants displayed fewer stress signals - suggesting a decrease in stress with continued exposure to training. Our findings align well with prior research showing similar patterns across multiple non-human species. Further, our findings suggest that voluntary cooperation can be achieved even under challenging circumstances and provide learning opportunities for both elephants and mahouts. Our work outlines a pathway to reshape captive animal welfare measures in India.

## 1. Introduction

In recent decades, animal training practices have undergone a significant transformation, shifting away from coercive and aversive techniques to more humane, evidence-based methods. Positive reinforcement training (PRT), which rewards desired behaviours with the addition of any stimulus pleasing to the subject, has gained widespread acceptance for its effectiveness and welfare benefits, especially in zoos and aquaria [36]. Spiezio[37] further found that obtaining voluntary cooperation in husbandry and veterinary procedures not only facilitates management but also improves welfare by reducing fear and providing cognitive stimulation. Most documented PRT initiatives have occurred in controlled environments such as zoos or sanctuaries [24,26].

Studies consistently demonstrate that punishment-based approaches are associated with increased stress and compromised welfare. For example, dogs trained with aversive techniques exhibit significantly higher stress levels than those trained using reward-based methods [29]. A comprehensive review by Ziv[38], covering 17 studies, concluded that the use of aversive methods can jeopardise both the physical and mental health of dogs.

PRT has been applied across a wide range of zoo-housed species. In primates, voluntary medical behaviours such as blood draws are now standard and are associated with decreased stress and increased prosocial behaviours [39]. In large carnivores, PRT enables animals to move between enclosures using recall cues and supports medical care without the need for sedation. Similarly, smaller-bodied mammals, such as meerkats, have been trained to voluntarily enter crates for veterinary separation [36]. The same review shows that even a Nile crocodile was station-trained for blood draws and weighing without sedation, and a gharial was trained to voluntarily enter a crate for transport.

Some regions in South Asia are beginning to adopt these methods. In Nepal, for example, trainers in zoos increasingly rely on PRT to encourage voluntary participation in husbandry and veterinary procedures, resulting in improvements in behaviour, management, and welfare [40]. Fagen et al.[25] reported that, in Nepal, PRT increased voluntary participation in trunk-wash procedures for tuberculosis screening from 39% to 89% within just 35 sessions.

However, the training of captive elephants in South Asia, particularly in India, continues to largely rely on traditional dominance-based methods. These practices include the use of tools such as the ankus (bullhook), physical restraint, and aversive conditioning that rely on fear or pain to elicit compliance [40]. Some of the documented impacts include physical injuries, psychological distress, and behavioural abnormalities [23,27].

The application of PRT to working elephants in India remains rare. Captive working elephants in India - which are used in logging, tourism, religious festivals, and patrolling, are particularly subject to these approaches. This study addresses the critical gap between established welfare-based training methods and current practices in India’s working elephant systems. It explores whether PRT can be used as a low-stress, welfare-positive training method for captive working elephants in India.

## 2. Materials and Methods

### 2.1. Study Area

This study was conducted in **Pakke Tiger Reserve** located in **Pakke Kessang District**, Arunachal Pradesh, Northeast India (26O 86’ N, 92O 84’ E). Covering approximately 862 km^2^, PTR lies in the Eastern Himalayan foothills and forms part of the larger Kameng Protected Area Complex. The landscape is characterised by semi-evergreen and tropical moist deciduous forests [1], steep terrain, and high annual rainfall, particularly during the monsoon season (May to September), when access to the reserve becomes extremely difficult.

PTR relies on **captive elephants for patrolling and conservation activities**. These elephants are stationed at various anti-poaching camps and serve critical roles in navigating challenging terrain, transporting rations, carrying out rescue efforts, and enforcing protection in areas where vehicular access is limited or impossible. However, these approaches have contributed to accidents, trauma, and deteriorating relationships between elephants and their handlers (mahouts), with recent years witnessing the deaths of two mahouts due to elephant attacks and the loss of two elephants to health complications.

### 2.2. Data Collection

We conducted the program from October 2024 to April 2025. A kraal was built at the Khari anti-poaching camp in Pakke Tiger Reserve prior to the program. The kraal (with four vertical corners each 4 meters in height) was secured (with 1 meter embedded in the ground and 3 meters above ground) and had nine horizontal wall posts fixed at three levels (60, 175, and 260 cm from the ground) to form three visible rails on each side.

The training was conducted at two anti-poaching camps - **Khari and Diji**, involving a total of **five elephants and eleven mahouts**. The program focused on encouraging voluntary participation of elephants in key medical behaviours (e.g., foot care, injections, blood draws), while building mahout capacity in PRT methods. We were unable to complete the program in Diji, so the analysis here only includes data from the three elephants in Khari. Table 1 shows the list of trained and desired behaviours.

**Table 1.** Progress of voluntary medical behaviour training in captive elephants at Pakke Tiger Reserve (Oct 2024 – Mar 2025): A comparative overview of completed, ongoing, and upcoming behaviours for Khaisingh, Bahadur and Vijaya.

| Behaviours | Khaisingh<br>(M, 31) | Bahadur<br>(M, 38) | Vijaya<br>(F, 48) |
| --- | --- | --- | --- |
| Front Foot Present (Right) | ✓ | ✓ | ✓ |
| Front Foot Present (Left) | ✓ | ✓ | ✓ |
| Back Foot Present (Right) | ✓ | ✓ | Need to train |
| Back Foot Present (Left) | ✓ | ✓ | Need to train |
| Side Line Up (Right) | ✓ | ✓ | Being trained |
| Side Line Up (Left) | ✓ | ✓ | Need to train |
| Back Foot Side Present (Right) | ✓ | ✓ | Need to train |
| Back Foot Side Present (Left) | ✓ | ✓ | Need to train |
| Ear Present/Voluntary Blood Draw (Right and Left) | ✓ | ✓ | Need to train |
| Injection: Shoulder (Right) | ✓ | Being trained | Need to train |
| Injection: Shoulder (Left) | ✓ | Being trained | Need to train |
| Temperature | ✓ | Being trained | Need to train |
| Eye Medication (Right) | ✓ | Being trained | Need to train |
| Eye Medication (Left) | ✓ | Being trained | Need to train |

We conducted two training sessions with the elephants every day - one in the morning, and the second in the afternoon. Each session was recorded either on a GoPro or an Android phone camera. The video length varied from 40 seconds to 20 minutes.

Our three elephants (aged 31-48) and their primary caregivers had spent ten or more years together. Bahadur, a 38-year-old male elephant with 20 years at Pakke, was primarily cared for by Debaru Guawala (18 years of experience) and Bikas Guawala, who has 15 years of experience and had worked with Bahadur for a year. Vijaya, a 48-year-old female elephant, has spent her entire life in the reserve and was primarily cared for by Jully Welly, who had worked with her for 14 years. He was supported by Chunumunu Guawala, who had around 13 years of experience.

Khaisingh, a 31-year old male elephant with 17 years in Pakke, was primarily cared for by Sangkhang Boro with 10 years of experience with elephants (including 2 years with Khaisingh). He was supported by Sonu Kharia, who had worked with Khaisingh for about 2 years. The training team was further supported by Rasham Brah, the head mahout at Khari, who has more than 25 years of experience with elephants.

The training sessions were designed and implemented by Ms. Chrissy Pratt, an animal trainer and behaviourist with over 20 years of experience in positive reinforcement training (PRT). At Pakke Tiger Reserve, she conducted approximately 170 training sessions with the three study elephants, contributing almost 20 hours of direct training. Her role included designing the training sessions, leading implementation in collaboration with mahouts, and mentoring them to continue sessions independently. Chrissy has previously applied similar methods at multiple international elephant facilities, including Save Elephant Foundation and Boon Lott’s Elephant Sanctuary (Thailand), Wild is Life (Zimbabwe), Tiger Tops Elephant Camp (Nepal), and the Myanmar Timber Enterprise, providing a strong foundation for adapting PRT to the socio-cultural context of Pakke.

### 2.3. Data Analysis

We decided on a list of behaviours we wanted to code in consultation with the elephant trainer, The Elephant Ethogram [6], and related research on African and Asian elephants as well as other animal species [7–16]. Table 2 lists the behaviours we decided on. Low stress is defined as low levels of avoidance and displacement behaviours, reductions in behavioural indicators of distress, and increased voluntary participation in husbandry routines.

**Table 2.**
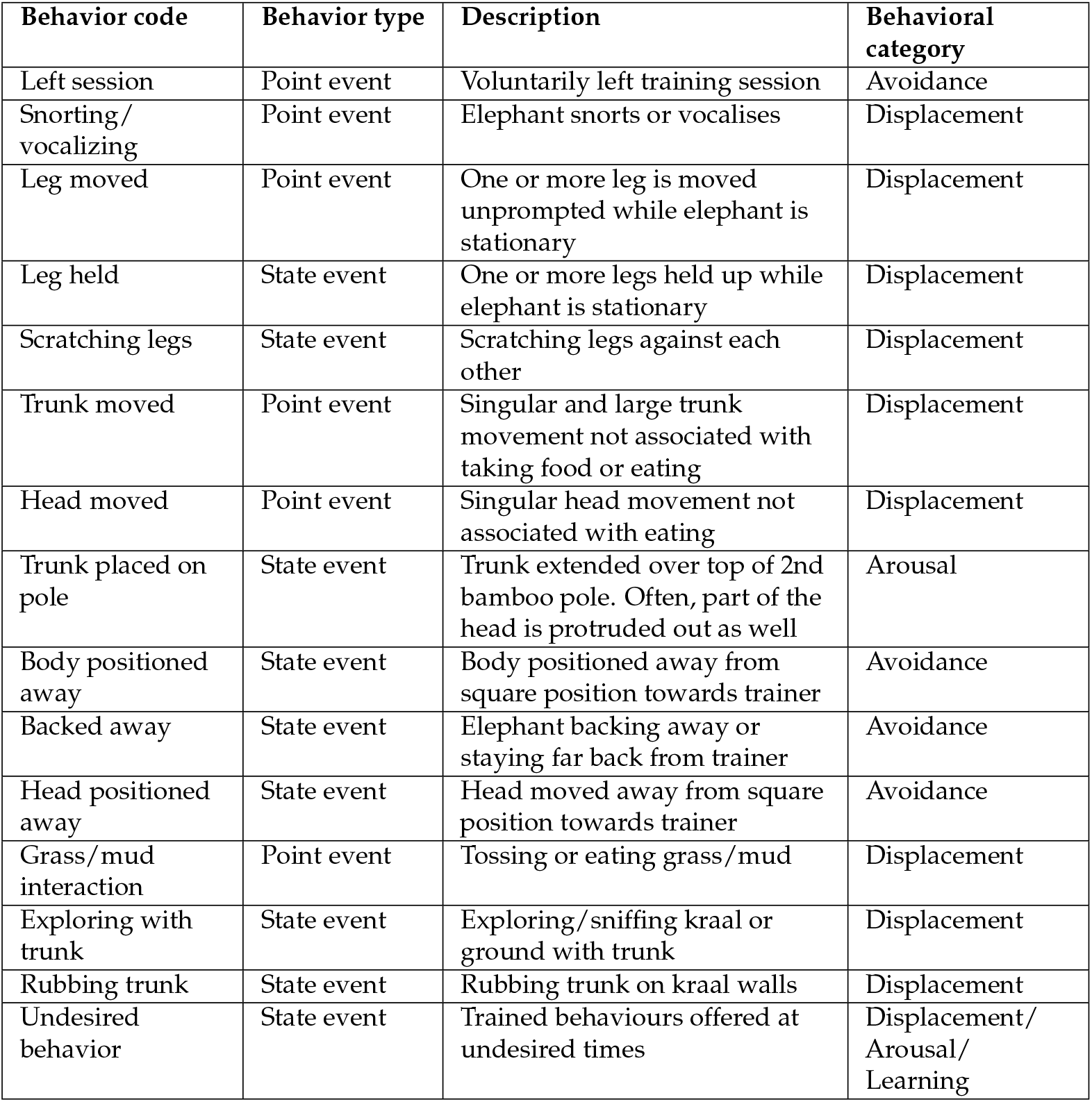
Ethogram of behaviors recorded during training sessions with captive elephants at Pakke Tiger Reserve.

| Behavior code | Behavior type | Description | Behavioral category |
| --- | --- | --- | --- |
| Left session | Point event | Voluntarily left training session | Avoidance |
| Snorting/vocalizing | Point event | Elephant snorts or vocalises | Displacement |
| Leg moved | Point event | One or more leg is moved unprompted while elephant is stationary | Displacement |
| Leg held | State event | One or more legs held up while elephant is stationary | Displacement |
| Scratching legs | State event | Scratching legs against each other | Displacement |
| Trunk moved | Point event | Singular and large trunk movement not associated with taking food or eating | Displacement |
| Head moved | Point event | Singular head movement not associated with eating | Displacement |
| Trunk placed on pole | State event | Trunk extended over top of 2nd bamboo pole. Often, part of the head is protruded out as well | Arousal |
| Body positioned away | State event | Body positioned away from square position towards trainer | Avoidance |
| Backed away | State event | Elephant backing away or staying far back from trainer | Avoidance |
| Head positioned away | State event | Head moved away from square position towards trainer | Avoidance |
| Grass/mud interaction | Point event | Tossing or eating grass/mud | Displacement |
| Exploring with trunk | State event | Exploring/sniffing kraal or ground with trunk | Displacement |
| Rubbing trunk | State event | Rubbing trunk on kraal walls | Displacement |
| Undesired behavior | State event | Trained behaviours offered at undesired times | Displacement/Arousal/Learning |

The behaviours were classified into three different categories for ease of analysis and visualisation. The categories we used are:

#### Avoidance

Avoidance refers to behavioural responses that function to increase distance from or reduce contact with aversive stimuli, potential threats, or stressful situations. These behaviours serve to minimise exposure to negative experiences and can include physical withdrawal, gaze aversion, or active movement away from the source of distress [17,20].

#### Displacement

Displacement behaviours are activities that appear out of context or are seemingly irrelevant to the immediate situation, typically occurring when an animal experiences conflicting motivations or stress. These behaviours often involve normal maintenance activities (such as grooming, feeding, or comfort movements) that occur at inappropriate times or in inappropriate contexts, suggesting that they serve as outlets for redirected energy or tension[12,19].

#### Arousal

Arousal encompasses the general activation state of an organism’s nervous system, reflecting the intensity of physiological and behavioural responses to stimuli. High arousal states are characterised by increased alertness, heightened responsiveness, and often elevated activity levels, while low arousal states involve reduced responsiveness and activity [18].

Videos were coded as per this framework using the BORIS software [2]. All videos from the first six weeks of training were coded. Not seeing significant changes in the presentation of stress signals thereafter, we then coded every session in which new training was introduced, and also made sure to code at least 5 sessions for each elephant picked randomly from every 7 days. We coded approximately 36 hours of video footage from October 2024 to March 2025, with a break in mid-December to mid-January (Khaisingh - 108 sessions, 15.2 hours, Bahadur - 107 sessions, 13.3 hours, Vijaya - 104 sessions, 7.7 hours).

Two of the behaviours listed in Table 2 were excluded from our analysis - *Leg held*, and *Undesired behaviour*. In teaching the elephants to put their feet up for examination and care, they were being asked to hold their legs up as an intermediate behaviour. Leg held was thus unreliable as an indicator of displacement during training the behaviour, so we decided to exclude it entirely.

*Undesired behaviour* seemed to indicate different categories in different contexts. While learning different behaviours, the elephants might not have known what was being asked of them, so they might have been offering different behaviours as a way to get their treats faster. Bahadur appeared to be using behaviours he was comfortable with as a way of avoiding behaviours he was not as accustomed to. Determining the category for each incidence of the behaviour would have become very subjective, so undesired behaviours were excluded from our category-level analyses as well.

All analysis was done using the Julia language [3]. All data-wrangling and summarization was performed using the DataFrames [4] and DataFramesMeta libraries. Plots were created using the Plots [5] and StatsPlots libraries.

While avoidance and displacement are clearly related to stress, elephants could exhibit high arousal states for various reasons - including anticipation of food, frustration, or being over-stimulated by the training context. Since high arousal might not necessarily be related to stress, we did not include it in our analyses. We used descriptive and summary statistics to study the relationship between stress signals and time - with the expectation that these signals would decrease over time. All correlation coefficients presented in the following sections are Spearman’s rank correlation coefficients. The non-linear relationship and expectation of monotonicity make this an ideal metric for capturing the strength of the relationship.

For averages across all three elephants, confidence intervals are not included due to the small sample size.

## 3. Results

Although our sample size was only three elephants, we found that the average of the stress signals across subjects decreased as sessions progressed, although the trend was not strictly monotonic (Figure 1). Correlation coefficients for all categories were less than -0.55, supporting our hypothesis that stress signals decrease as the elephants participate in more sessions (Table 3). p-values for most of the categories showed that the correlation was significant. The only category with a p-value close to 0.05 had one behaviour in it (Left session), was presented by only one elephant (Vijaya), and only 7 times. Table 3 shows that all of our categories have a strong negative relationship with session number-showing that the stress signals we coded reduced as the elephants participated in more sessions.

**Table 3.** Correlation coefficients for the different categories of behaviours. High p-values lend credence to the null hypothesis - that there is no relation between stress signals and session number, while a low p-value suggests that the null hypothesis is likely to be untrue. The t_statistic is a measure of the strength of the relationship. Values greater than 2 are considered significant, while values greater than 3 indicate strong relationships.

| Category | Correlation | p_value | t_stat |
| --- | --- | --- | --- |
| Avoidance (state) | -0.67 | 0.00 | -9.14 |
| Displacement (state) | -0.61 | 0.00 | -6.06 |
| Avoidance (point) | -0.78 | 0.04 | -2.84 |
| Displacement (point) | -0.58 | 0.00 | -7.24 |

**Figure 1.**
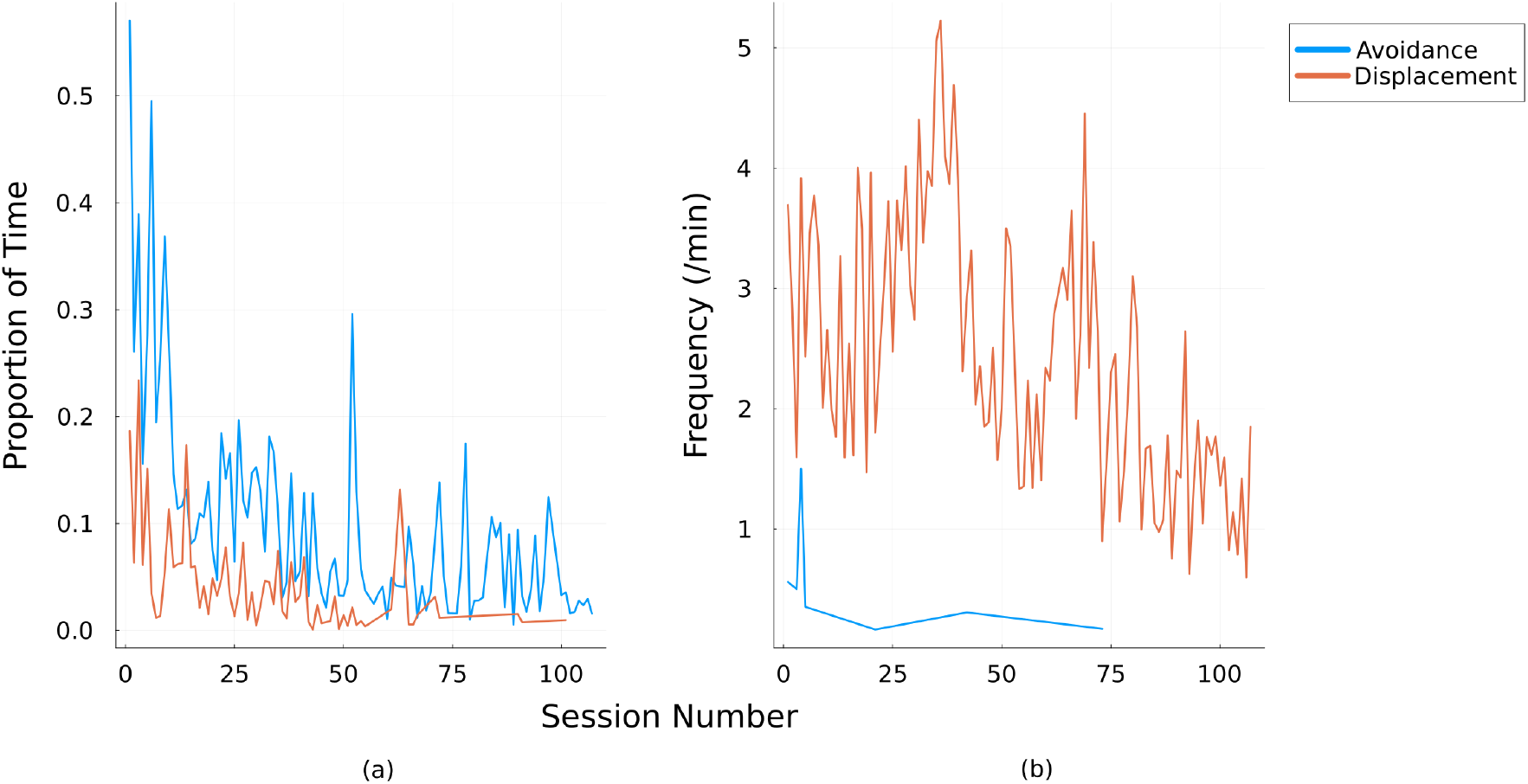
Average of stress indicators classified into categories as sessions progress. (a) shows state behaviours, while (b) shows point behaviours.

These trends of decreasing stress signals over time held true at the individual level as well, although there were differences between each of the elephants (Figures 2, 3, A1, A2). For example, Bahadur consistently displayed higher durations of avoidance behaviours (such as backing away from the trainer) than the other two elephants throughout training.

**Figure 2.**
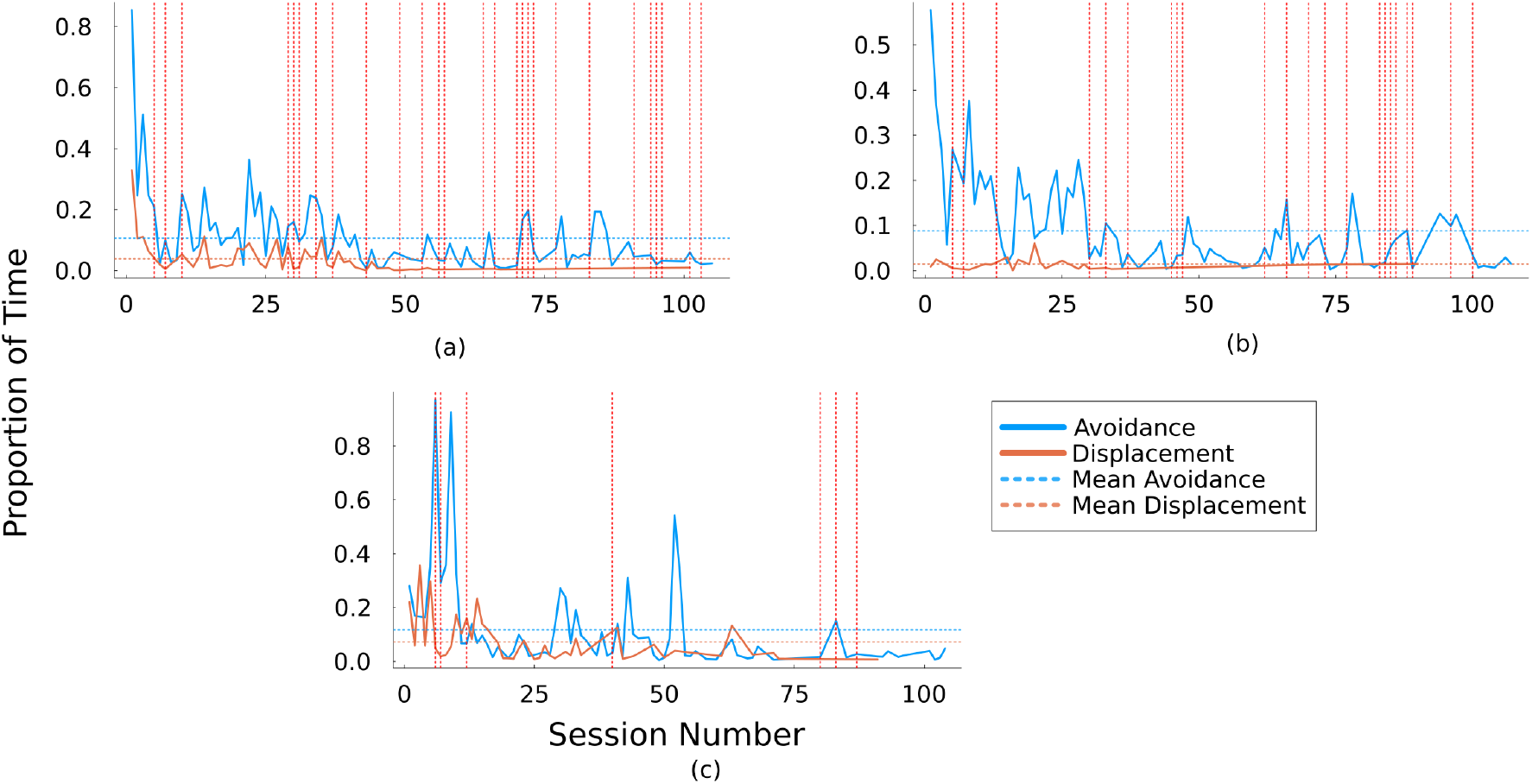
The trajectory of *state behaviours* over time grouped into categories for (a) Khaisingh, (b) Bahadur, and (c) Vijaya. The dashed horizontal lines show means for each category. Dashed vertical lines signify sessions when new behaviours were introduced.

**Figure 3.**
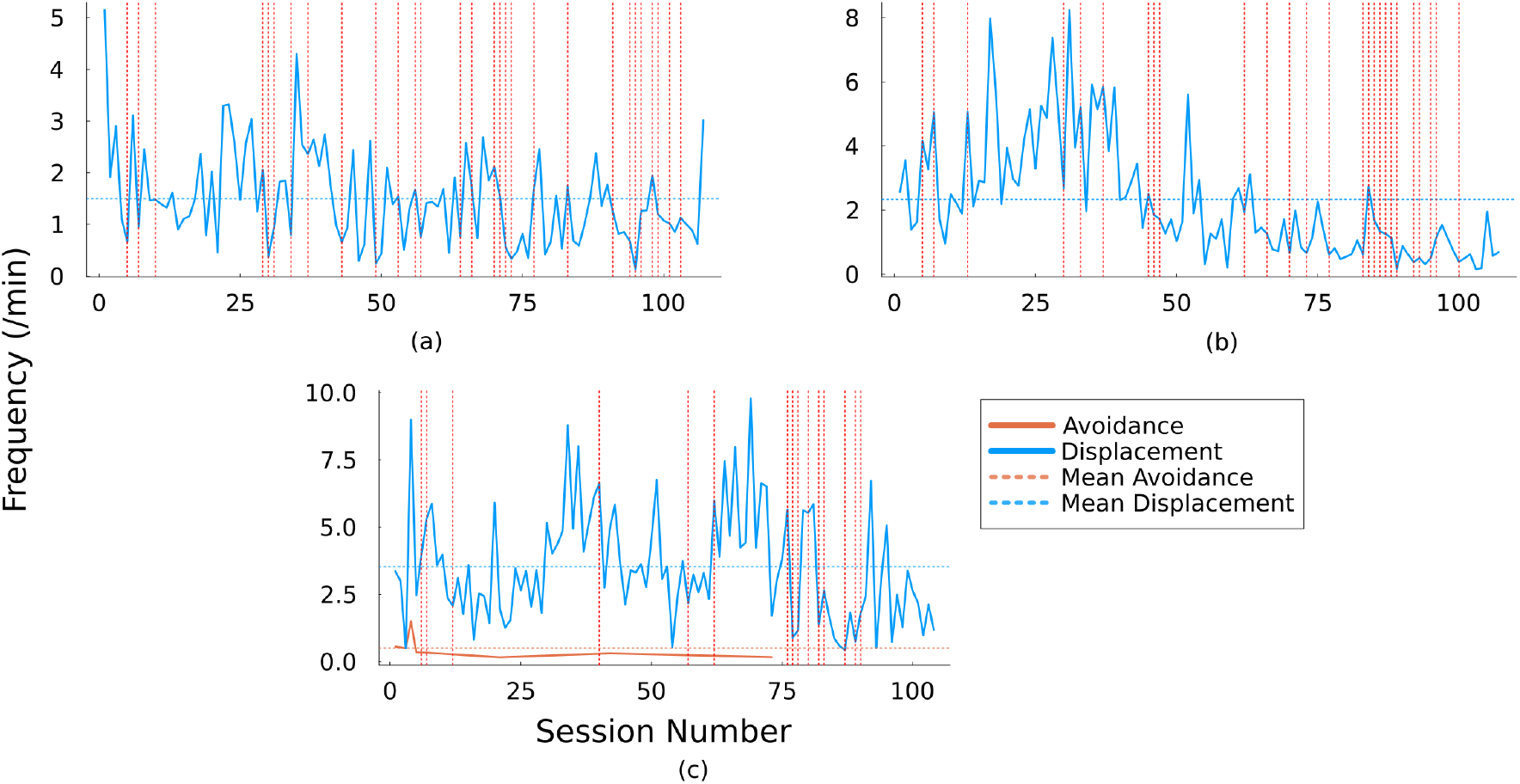
*Point behaviour* trend lines for (a) Khaisingh, (b) Bahadur, and (c) Vijaya grouped by category. Dashed horizontal lines show the means for categories. Dashed vertical lines demarcate sessions when new behaviours were introduced.

The correlation coefficients for avoidance for all three elephants were similar (Bahadur -0.56, Khaisingh -0.47, Vijaya -0.57 for state events, Vijaya - 0.78 for point events), showing that the avoidance category was negatively correlated with session number. Correlation coefficients and p values for all three (r = -0.48, p-value 0 for Vijaya (state), r = -0.61, p-value 0 for Khaisingh (state), and r = -0.71, p-value 0 for Bahadur(point)) provided strong support for displacement events reducing as sessions progressed as well. Vijaya left the training session a few times; which accounts for the presence of the avoidance category in her point events graph (Figure 3).

While there were some differences between state and point events and between individuals, the results of our analysis validated our expectation of stress signals decreasing as the elephants participated in more sessions.

## 4. Discussion

Our goal in conducting this analysis was to determine whether stress signals reduced as session number increased. All of our results and observations seem to be in line with prior findings. Physiological stress reduction during repeated positive training experiences has been documented in other species. In shelter dogs participating in animal-assisted interventions, salivary cortisol levels showed a significant decrease from baseline measurements at the beginning compared to the end of an 8-week training program, suggesting that repeated positive training experiences led to long-term reductions in physiological stress indicators [28]. This aligns well with our primary finding about stress signals decreasing with regular, recurring training sessions. Few studies have documented changes in behaviour patterns during training. Our study thus fills an important gap in the longitudinal documentation and characterization of stress signals in the course of a planned training intervention.

Our behavioural observations of decreasing displacement and avoidance behaviours over successive training sessions align with the observed pattern of stress reduction through positive training experiences. Research on cattle has shown that habituation protocols can improve behavioural responses over time, with animals showing “improved numerical behavioural scores” and decreased cortisol concentrations from the start of experiments to 14 days after treatment initiation, demonstrating that behavioural adaptation to handling can occur with repeated positive exposure [35].

Individual variation in training responses has also been well-documented across species. In rhesus macaques, research has shown significant individual differences in training success, with animals categorized as “exploratory” (over 85% successful), “moderate” (over 75% successful), or “inhibited” (approximately 22% successful) in learning target-touching tasks, demonstrating that temperament affects training responsiveness [30]. In our project as well, all three elephants had different responses to training. Vijaya seemed to be much slower at picking up new behaviours (Table 1) and did not take to new people. She would leave training sessions and showed more tendencies towards aggression than either of the bulls, so we kept the rate of reinforcement high and the training short to ensure that the sessions were rewarding for her.

Studies in fish have similarly found that personality traits significantly affect learning ability, with shy individuals being more successful at spatial learning tasks compared to bold individuals, suggesting widespread effects of individual characteristics on training outcomes across species [31]. Khaisingh was extremely food motivated (at times too motivated) and would offer undesired behaviours if the trainer did not deliver rewards fast enough, so we also had to try and communicate that simply standing still and not doing anything was a desired action as well. New behaviours appeared to stress Bahadur resulting in avoidance behaviours or offering familiar habituated behaviours.

We have anecdotal findings that when new behaviours were introduced, our elephants would regress to displaying higher levels of stress signals (Figures 2 and 3 ridges at behavioural milestones). The concept that stress signals return when novel behaviours are introduced aligns with established principles of learning and habituation. Research has shown that “habituation can be disrupted by almost any change in the experimental conditions” and that “the presentation of a novel stimulus may restore the response on the next trial,” a phenomenon known as dishabituation [34]. Displacement behaviours are believed to occur when animals face conflict between competing motivations, and stress can return when animals encounter novel situations or tasks that create new conflicts.

## 5. Conclusions

Future iterations of this work could include conducting comparisons of stress signals before and after elephants have been taught to voluntarily participate in husbandry. Studies have documented behavioural differences between training methods - with dogs trained using aversive methods displaying significantly more stress-related behaviours including avoidance behaviours such as body turns, moving away, crouching, and lying on side/back, compared to dogs trained with positive reinforcement [29]. Demonstrating this pattern in working elephants would be an important step towards the wider adoption of low-stress handling methods.

We would also like to measure the efficacy of our training. One of the objections raised against positive reinforcement based training is that it takes much longer than traditional methods. More training programs and proper documentation would enable us to learn how best to convey these methods to mahouts and determine personality traits that make particular elephants more or less amenable to the training. We strongly believe that the resultant safety for elephants, mahouts, and veterinarians and the improved relationships between elephants and mahouts far outweigh the time taken to get results. Obtaining the voluntary cooperation and continued training of handlers might have positive implications on animal welfare through challenging circumstances and learning opportunities.

## Author Contributions

“Conceptualization, R.K. and C.P.; methodology, R.K. and C.P.; validation, S.R, P.R. and N.V.; formal analysis, R.K.; resources, K.R., S.R., N.V. and P.R.; data curation, R.K. and A.B.; investigation, C.P., S.R., P.R., R.K. and A.B.; software, R.K.; visualization, R.K.; writing—original draft preparation, N.V., R.K. and S.R.; writing—review and editing, N.V., R.K. and S.R.; supervision, N.V.; project administration, S.R., P.R. and K.R.; funding acquisition, S.R. and P.R.

## Data Availability Statement

The original data presented in the study are openly available in Github at the linked repository.

## Acknowledgments

This study was made possible through an official collaboration between the Pakke Tiger Reserve (PTR) Forest Department and the Pachyderm Partnerships Project of the Canopy Collective, formalised in October 2024. We are deeply grateful to the PTR Forest Department for their permissions, logistical support, and for providing a space to build the training kraal. Our heartfelt thanks go to the mahouts and jugalis of Pakke, whose experience, care, and commitment made this work possible: Rasam Brah, Debaru Guawala, Bikas Guawala, Juli Welly, Sankhang Boro, Chunumunu Guawala, Sonu Kharia, Pali Nabum, Brajen Saikia, Kaka Tallang, Lucky Nabum, and Jiskal Nabum. We are especially thankful to the elephants Khaisingh, Bahadur, Vijaya, Joimala, and Tamuk, who were not just participants but teachers in their own right. We also thank our advisors, Dr. Abhilasha Sharma and Dr. Rinku Gohain, for their encouragement and guidance. We are grateful to our funders, Rohini Nilekani Philanthropies and The Habitats Trust, for believing in this work; and to the Dusty Foot Foundation and the Seegreen Foundation for logistics and administrative support. Finally, we would like to acknowledge the Forest Department staff Satyaprakash Singh, IFS (DFO, PTR), Singku Maga (RFO, Seijosa) and Geto Marde (Forester, Seijosa) and most especially the staff at Khari and Diji, Montu Basumatari, Gangamuni Guawala, Rajo Tayem and Limar Loyi, who made the camps a true home away from home.

## Abbreviations

The following abbreviations are used in this manuscript:

PTR: Pakke Tiger Reserve
PRT: Positive Reinforcement Training

## Appendix

**Figure A1.**
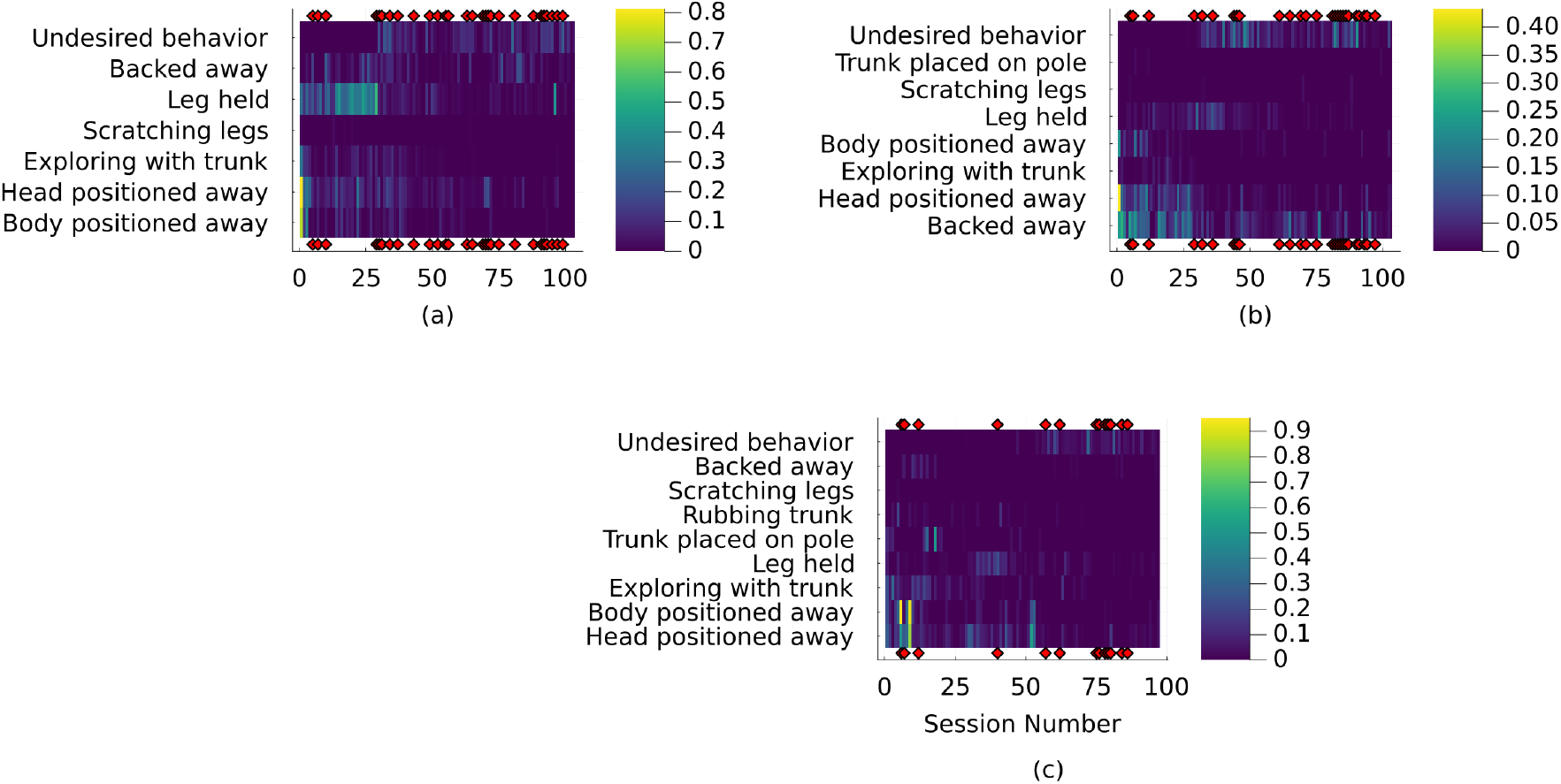
Heatmaps of time budgets for *state events* for all three elephants. The colour represents proportion of the session spent displaying the behaviour.

**Figure A2.**
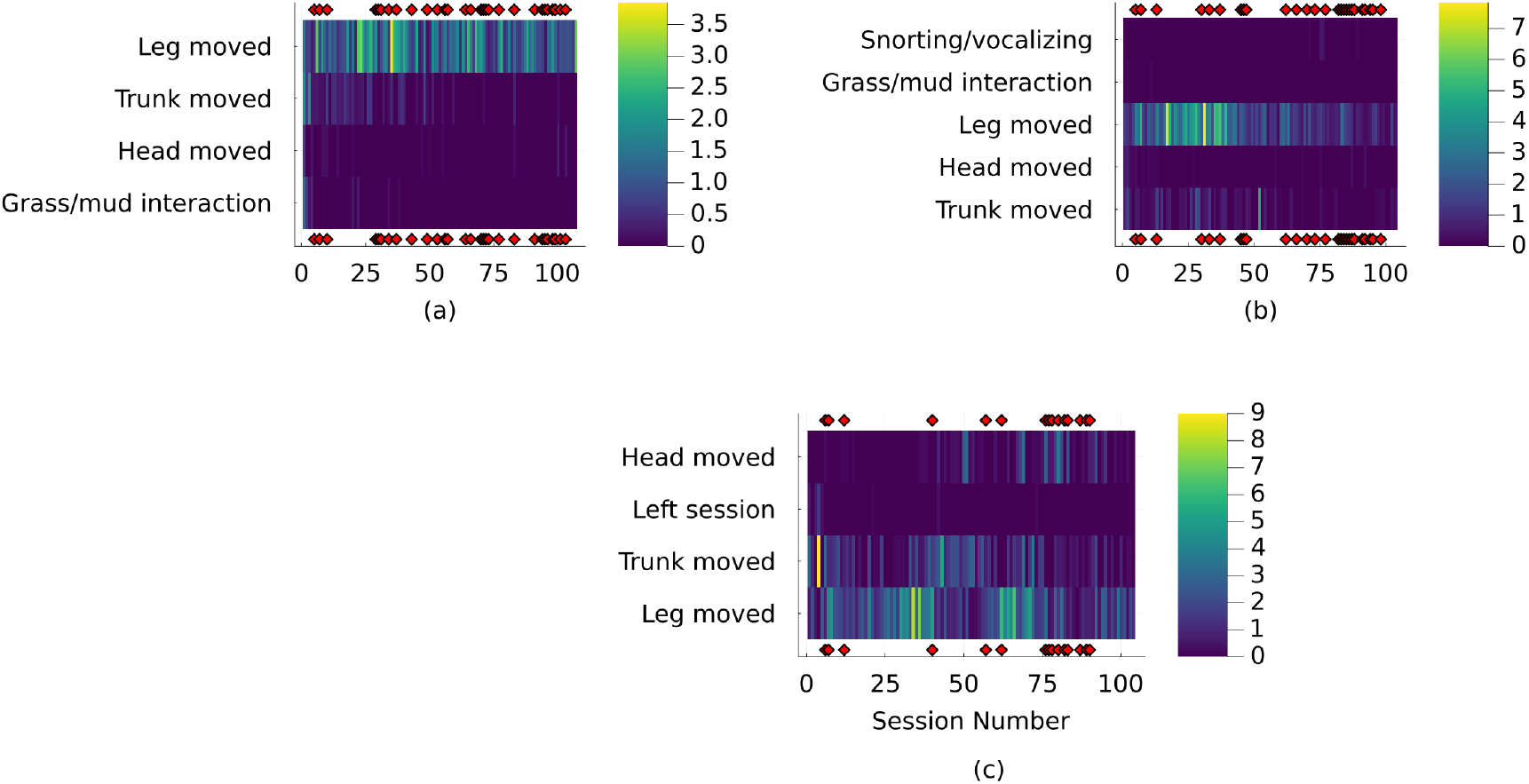
Heatmaps of behaviour frequencies for point events for all three elephants. The colour represents frequency (/min) of the behaviour

## Disclaimer/Publisher’s Note

The statements, opinions and data contained in all publications are solely those of the individual author(s) and contributor(s) and not of MDPI and/or the editor(s). MDPI and/or the editor(s) disclaim responsibility for any injury to people or property resulting from any ideas, methods, instructions or products referred to in the content.

